# Hormonal and transcriptomic analysis reveals the role of ABA and BR in breaking the barrier of inter-subgeneric hybridization in water lily (*Nymphaea*)

**DOI:** 10.1101/2023.09.05.556322

**Authors:** Ping Zhou, Jingwen Li, Huiyan Jiang, Zhijuan Yang, Chunqing Sun, Hongyan Wang, Qun Su, Qijiang Jin, Yanjie Wang, Yingchun Xu

**Author notes:** Corresponding authors. E-mail addresses (Yingchun Xu).

## Abstract

Understanding the process of signal communication between pollen and stigma is of significant importance for plant sexual reproduction. In the case of inter-subgeneric hybridization in water lily, there exists a pre-fertilization hybridization barrier, the regulatory mechanism of which remains unclear. In this study, we conducted hormone and transcriptome analyses of unpollinated stigmas (Mock), self-pollinated stigmas (SP), cross-pollinated stigmas within the same subgenus (CP), and inter-subgenus cross-pollination stigmas (ISCP) in water lily to elucidate the formation mechanism of the inter-subgeneric hybridization barrier. Our results indicated that the lack of ABA and BR in ISCP stigmas are key factors contributing to the formation of the inter-subgeneric hybridization barrier in water lily. Exogenous application of ABA and BR can help overcome the barrier between inter-subgeneric water lily crosses. Through transcriptome analysis, we identified nine candidate genes involved in regulating the inter-subgeneric hybridization barrier in water lily. In addition, we further demonstrated the importance of the NCED2-mediated ABA synthesis pathway in the hybridization process through AS-ODN technology. Our study confirms that ABA and BR are critical for breaking the inter-subgeneric hybridization barrier in water lily. The identification of the nine candidate genes provides important clues for further research on the hybridization recognition mechanism in water lily.

## Introduction

Water lily is a collective name for plants of the *Nymphaea*, which are widely used in landscaping, medicine and water restoration (Parveen *et al*., 2022; Xiong *et al*., 2023; Yu *et al*., 2018). To date, more than 50 species of water lilies have been discovered, which can be categorized into tropical and hardy water lily according to their geographical distribution (Perry and Robinson, 1997; Sun *et al*., 2021). Tropical water lily is favored for their diverse flower colors, including red, blue, purple, white, yellow and other complex hues (Yu *et al*., 2018). However, tropical water lily is intolerant to low temperatures and struggle to survive harsh winters in higher latitudes (Hao *et al*., 2022). On the other hand, hardy water lily, found in temperate regions, possess unique cold tolerance (Sun *et al*., 2023; Sun *et al*., 2021). Nevertheless, the flower colors of hardy water lily are limited to red, white and yellow, and their flower stalks do not emerge above the water surface, reducing their ornamental value (Sun *et al*., 2023; Sun *et al*., 2021). Therefore, hybrid breeding between tropical and hardy water lily holds great significance in cultivars that are both cold-tolerant and possess diverse flower colors. However, the hybridization between tropical and hardy water lily has proven to be challenging, with low seed set and inefficient pollination (Hao *et al*., 2022; Sun *et al*., 2019; Sun *et al*., 2018). Studies have shown that pre-fertilization hybridization barriers during the cross between tropical and hardy water lily are the main hurdles in achieving successful seed production (Hao *et al*., 2022; Sun *et al*., 2019; Sun *et al*., 2018), with low pollen germination rate being a commonly observed phenomenon. Additionally, abnormalities in pollen tube growth, such as bending or swelling at the tip and failure to penetrate the ovule, have been observed (Hao *et al*., 2022).

Studies have shown that indoleacetic acid (IAA), zeatin riboside (ZR) and abscisic acid (ABA) are the key hormones involved in plant fertilization (Breygina *et al*., 2023; Kovaleva *et al*., 2017; Mesejo *et al*., 2013). In pollen germination experiments, IAA and gibberellic acid (GA) stimulate pollen tube growth, while ABA inhibits pollen tube growth, and ZR has no significant effect on this process (Breygina *et al*., 2023). Low levels of IAA can lead to abnormalities in pollen tube growth (Ma *et al*., 2013). IAA levels positively correlated with the speed of fertilization, ABA promoting fertilization and ZR mainly related to sperm-egg fusion and seed development (Yang *et al*., 2022). In tobacco, IAA has been reported to participate in the interaction between pollen and stigma (Chen and Zhao, 2008). Jasmonic acid (JA) and salicylic acid (SA), closely related to biotic stress, have also been reported to participate in regulating plant pollination and fertilization processes (Lu *et al*., 2021; Schubert *et al*., 2019; Shi *et al*., 2017; Wang *et al*., 2021a). In pear, it has been reported that pollination can affect the synthesis of JA and ABA in the pistil. Pollination combinations with hybridization barriers activate the expression of genes related to phytohormone signaling pathways and plant-pathogen interaction pathways (Shi *et al*., 2017). It is believed that the JA-mediated defense pathway may be an important pathway leading to self-incompatibility in pears (Shi *et al*., 2017).

In this study, we examined changes in endogenous hormone contents in water lily stigmas after unpollinated (Mock), self-pollinated (SP), cross-pollinated in the same subgenus (CP) and inter-subgenus cross-pollination (ISCP) treatments. Changes in the contents of ABA, GA3, BR, MeJA and SAG were considered to be the main factors contributing to the formation of cross-subgenus hybridization barriers in water lily. Potential genes regulating the formation of cross-subgenus hybridization disorder in water lily were excavated by RNA-seq analysis. The role of *NCED2* in regulating the cross-subgenus hybridization disorder of water lily was further verified. The mining of this information provides a theoretical basis for understanding the mechanism of the formation of inter-subgenus hybridization barriers in water lily and improving the efficiency of inter-subgenus hybridization in water lily.

## Materials and methods

### Plant materials

This study used hardy water lily (*N.* ‘Peter Slocum’) and tropical water lily (*N.* ‘NangKwaug Fah’) as experimental materials. The water lily plants were preserved in the Bai Ma Education Demonstration Base of Nanjing Agricultural University. There are two main reasons for choosing these two cultivars for this study. Firstly, both cultivars have high fruit set rates and good pollen vitality, eliminating the influence of fruit set rates and pollen vitality on the experiment. Secondly, both cultivars exhibit obvious pre-fertilization hybridization barriers. *N.* ‘Peter Slocum’ and *N.* ‘NangKwaug Fah’ were used as the female and male parents, respectively. The hybridization was conducted following the methods described by (Sun *et al*., 2019; Sun *et al*., 2018).

### Phytohormone detection by LC-MS/MS

The stigma of water lily was collected after 6 h of pollination, including unpollinated (Mock), self-pollination (SP), cross-pollinated in the same subgenus (CP) and inter-subgenus cross-pollination (ISCP) treatments. Each sample had 4 biological replicates. After being flash frozen in liquid nitrogen, the samples were stored in a −80°C freezer. The levels of ABA, IAA, GA, JA, SA and brassinolide (BR), which belong to the six major classes of hormones, were detected in the stigmas of water lily.

The stigmas were frozen in liquid nitrogen, ground into fine powder, and 50 mg of powder was taken for extraction in a mixture of methanol: water: formic acid (15:4:1, v/v/v). The extraction solution was filtered through a 0.22 μm PTFE membrane (Anpel, Shanghai, China) and then analyzed using LC-ESI-MS/MS (Hui *et al*., 2018). The obtained mass spectrometry data were qualitatively analyzed by establishing the MWDB plant hormone database based on relevant standards (Šimura *et al*., 2018). The hormone content detection experiment was conducted at Suzhou Panomic Bio-medical Technology Co., Ltd. (Suzhou, China).

### Hormone treatments and pollen tube visualization

Removed the stamens of the first day blooming waterlily flower, and then add 30 uM ABA, 0.3 mM GA or 0.3 mg/L BR into the pistil fluid of the waterlily flower. After about 10 minutes of exogenous hormone application, perform pollination.

After 6 hours of pollination, fixed the stigmas in FAA fixative for 6 hours, then soften them overnight in 1 mM NaOH. Stained the softened stigmas with aniline blue using the experimental protocol described in (Duan *et al*., 2014; Su *et al*., 2020; Zhang *et al*., 2021). After making temporary slides, observed them under a fluorescence microscope at a wavelength of 405 nm and capture images using a DS-Ri2 digital camera.

### Antioxidant enzyme activity assay

For the determination of antioxidant enzyme activity, 1 mL of ice-cold 25 mM HEPES buffer (pH = 7.8) containing 0.2 mM EDTA, 2 mM ascorbic acid and 2% polyvinylpyrrolidone was used to extract total protein from 0.1 g of waterlily stigmas. The homogenate was centrifuged at 12,000g for 20 mins at 4°C, and then the supernatant was carefully collected for enzyme activity assays. The enzyme activities of superoxide dismutase (SOD), peroxidase (POD) and catalase (CAT) were measured. The measurement of enzyme activity follows the method described by (Wang *et al*., 2011).

### RNA extraction and sequencing

The samples used in this experiment are the same as the samples described in the hormone detection. The FastPure Universal Plant Total RNA Isolation Kit (Novazyme Biotechnology, China) was used to extract total RNA from plants, following the instructions of the kit. The integrity of the total RNA was determined by 1% agarose gel electrophoresis. The integrity and concentration of the RNA were accurately measured using the Agilent 2100 RNA Nano 6000 Assay Kit (Agilent Technologies, CA, USA). The detection indicators included RIN value, 28S/18S ratio, presence of baseline uplift in the profile and 5S peak. RNA-seq was performed by Suzhou Panomic Bio-medical Technology Co., Ltd (Suzhou, China) on the Illumina Novaseq6000 platform.

### RNA sequencing data processing

After filtering adapter sequences and low-quality regions, the clean reads were aligned to the *N. colorata* genome using HISAT2 (Kim *et al*., 2015; Pertea *et al*., 2016). The gene expression levels for each sample were calculated using featureCounts (Liao *et al*., 2014), specifically as FPKM (fragments per kilobase of exon per million mapped fragments). An expression matrix was generated using Trinity to assess differential gene expression among different samples (Grabherr *et al*., 2011). To identify enriched differentially expressed genes (DEGs) in terms of GO terms and metabolic pathways, functional enrichment analyses, including GO and KEGG, were performed relative to the whole transcriptome background using a bonferroni-corrected P-value threshold of < 0.05. For GO functional enrichment and KEGG pathway analysis, the R package ClusterProfiler was utilized (Wu *et al*., 2021).

### Co-expression network analysis

We performed co-expression network analysis of hormones and gene expression using the weighted gene co-expression network analysis (WGCNA) R package (Langfelder and Horvath, 2008). To determine the relationship between gene expression and hormone levels, we analyzed the normalized gene expression data along with the levels of ABA, GA3, Me-JA, SAG, and BR. We visualized module correlations and co-expression networks using the LinkET R package and Cytoscape software (v3.9.1) (Huang, 2021; Smoot *et al*., 2011). The obtained genes were further analyzed for correlation with genes involved in hormone synthesis, and annotation of the final genes was performed using hmmscan. Based on the results of qRT-PCR, we selected the most feasible genes for follow experiments(Johnson *et al*., 2010).

### Quantitative real-time PCR

HiScript III RT SuperMix for qPCR (gDNA wiper) kit (Novazyme Biotechnology, China) was used to synthesize the first strand cDNA. 1 μg total RNA was used in each 20 μL reaction. The cDNA products were diluted 10 times with deionized water before use. qRT-PCR experiments were conducted on a CFX96 Touch™ Real-Time PCR detection system (Bio-Rad, USA) using the ChamQ SYBR qPCR Master Mix kit (Novazyme Biotechnology, China). Each reaction used 2 μL of diluted cDNA, and other reaction components and conditions were carried out according to the manufacturer’s instructions. Specific primers were designed for qRT-PCR (Table S4). The *Actin11* gene of waterlily was used as the internal reference gene (Luo *et al*., 2010), and three biological and three technical replicates were performed for each treatment. The relative expression levels were calculated using the 2^-ΔΔCt^ method (Livak and Schmittgen, 2001; Xu *et al*., 2012).

### Oligonucleotide design and treatment

Design positive (S-ODN) and antisense (AS-ODN) oligonucleotides based on the cDNA sequence of NCED2 using the Sfold database (http://sfold.wadsworth.org/cgi-bin/soligo.pl) (Li *et al*., 2022; Su *et al*., 2020; Zhang *et al*., 2021; Zhao *et al*., 2020)(Table S5). Assess potential off-target effects using the BLAST program (https://blast.ncbi.nlm.nih.gov/Blast.cgi). The 5’ and 3’ ends of S-ODN and AS-ODN were modified with thiol groups to maintain stability. S-ODN and AS-ODN were synthesized by Beijing Tsingke Biotech Co., Ltd (Beijing, China).

Before ODN treatment, remove the stamens from selected water lily flowers first. Then, the designed ODN (the secretions from water lily stigmas were diluted to a concentration of 20 μM) was added to the stigmas of waterlily flowers. After 1 h of ODN treatment, the pollination procedure was performed.

### Statistics

For the analysis of intergroup significant differences, one-way analysis of variance (ANOVA) was used. When *P* < *0.05*, Tukey’s test was employed for post-hoc comparisons. Two-group comparisons were conducted using a two-tailed Student’s t-test. All analyses were performed using GraphPad Prism 8 software.

## Results

### Hormones play an important role in pollen and stigma recognition in water lily

To understand the regulatory of plant hormones on pollination in water lily, we analyzed the hormone levels in the stigma (Supplementary Table S1). The results revealed a significant accumulation of ABA, GA3, jasmonic acid-isoleucine (JA-lle) and BR in stigmas treated with SP and CP. In contrast, the levels of these four hormones were lower in Mock and ISCP treatments (Fig. 1). Interestingly, the content of GA8 in the stigma exhibited an opposing trend to that of GA3. This could be attributed to the conversion of active gibberellins to inactive gibberellins in the stigma after Mock and ISCP treatments. Methyl jasmonate (MeJA) and glycosylated salicylic acid (SAG), which are closely associated with biotic stress, were only detected in stigmas treated with ISCP. This suggests that certain regulatory networks associated with disease resistance in water lily stigmas were activated after ISCP treatment (Fig. 1). In addition, other hormones such as IAA, SA and JA appeared to have no direct correlation with pollination treatments (Fig. 1; Supplementary Fig. S1; Supplementary Table S1).

**Fig. 1.**
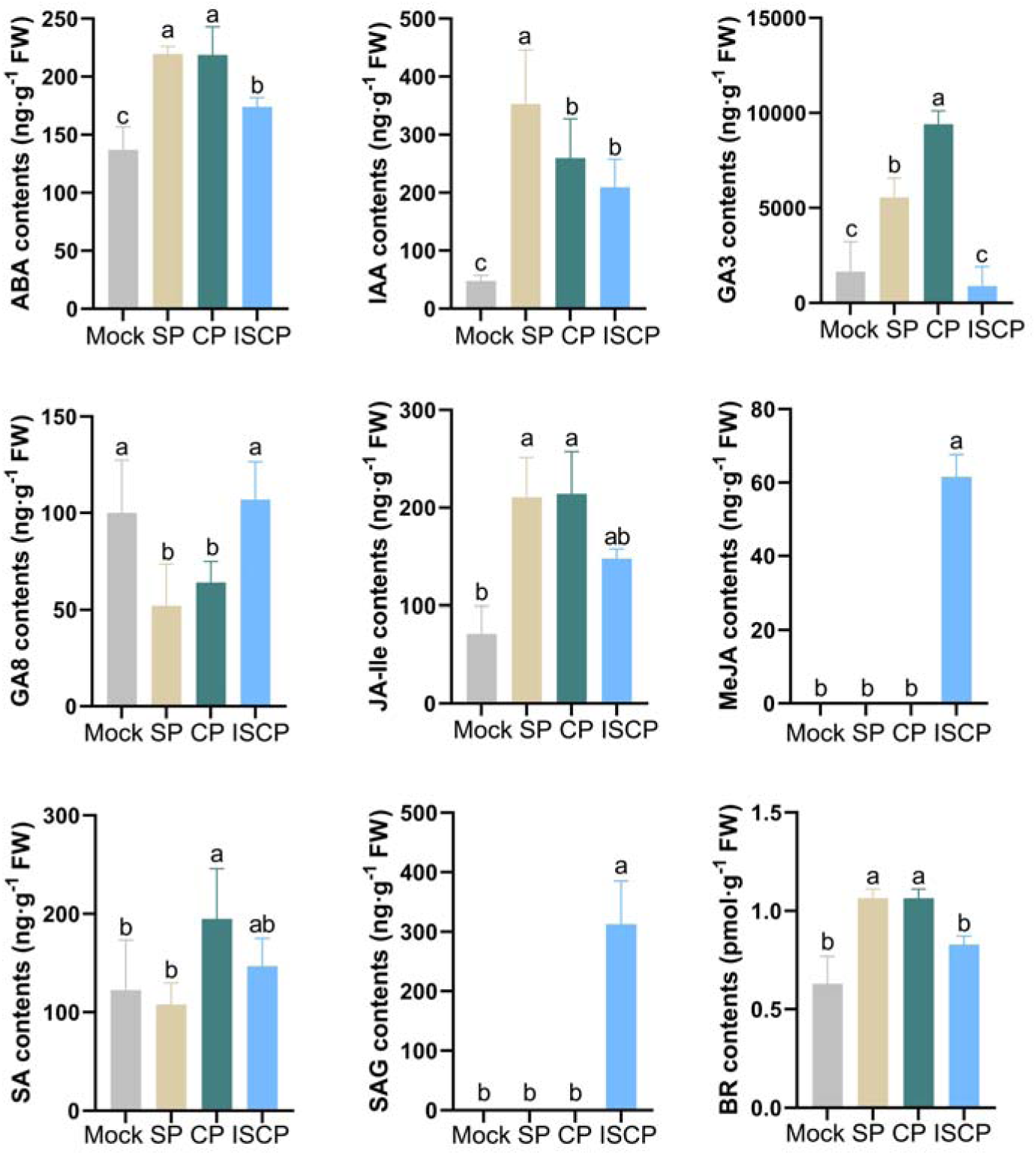
Analysis of phytohormones detected of Mock, SP, CP and ISCP stigmas in water lily. Concentrations of various hormones are expressed as mean ± SD. Different letters indicate statistically significant differences (*P* < *0.05*). Concentrations of other hormones are shown in Supplemental Fig S1 and Table S1.

### ABA and BR treatments increase fruit set after hybridization between water lily subgenera

To further investigate the role of hormones in the pollination hybridization process of water lily, we applied ABA, GA and BR onto the stigmas treated with ISCP. After a 6-hour treatment, we collected the pollinated stigmas and observed the germination of pollen tubes on the stigmas. The results showed a significant germination of pollen tubes on stigmas treated with SP and CP, while no pollen tube germination was observed on stigmas treated with ISCP (Fig. 2AB). When exogenous ABA and BR were applied, we observed that they promoted the germination of pollen tubes on ISCP-treated stigmas. However, GA treatment did not promote pollen tube germination on ISCP-treated stigmas (Fig. 2AB).

**Fig. 2.**
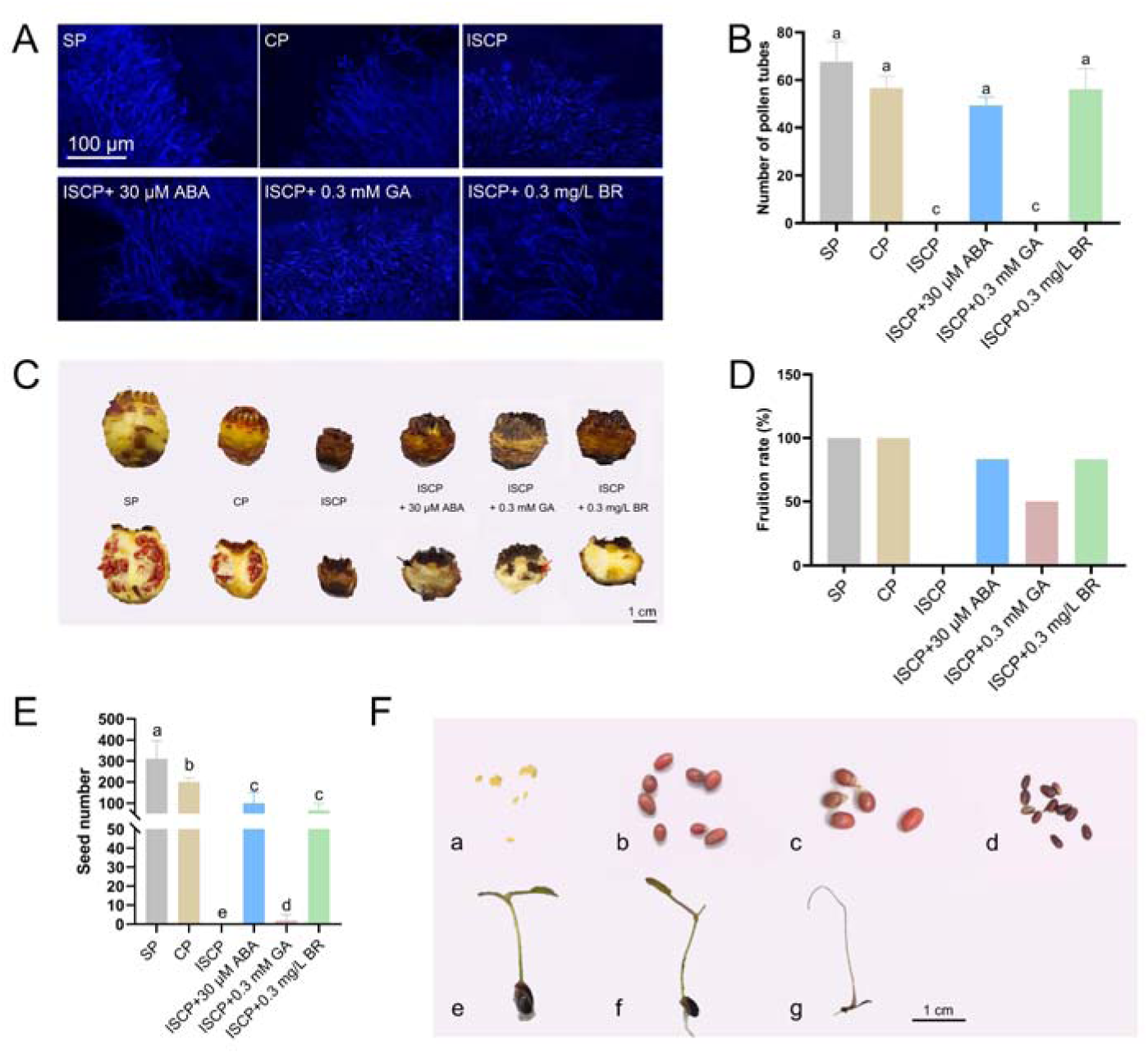
ABA and BR have the potential to break reproductive barriers of inter-subgenus in water lily. (A) Aniline blue staining showing the germination of pollen on water lily stigmas, Scale bar = 100 μm. (B) Statistics on the number of pollen germinating on water lily stigmas. (C) Effect of exogenous hormones on the fruiting of water lily (Black arrows indicate developing embryos, red arrows indicate abortive embryos) Scale bar = 1 cm. (D) Statistics on water lily fruiting rate (6 biological replicates per treatment). (E) Statistics on the number of seeds in fruits obtained under different treatments. (F) Seed morphology obtained after Mock, SP, CP, and ISCP treatments. (a) unpollinated embryos. (b) Mature seeds obtained with SP treatment. (c) Mature seeds obtained with CP treatment. (d) Mature seeds obtained with ISCP treatment (exogenous ABA and BR treatments). (e) Seedlings obtained with SP treatment. (f) Seedlings obtained with CP treatment. (g) Seedlings obtained with ISCP treatment. Scale bar = 1 cm.

To further understand the relationship between ABA, BR, GA and the fruit set rate in water lily hybridization, we collected the hybrid fruits of water lily after approximately 25 days of treatment. The results revealed that water lily treated with SP and CP successfully produced fruits with a high fruit set rate, and the seeds developed normally. However, water lily treated with ISCP did not show fruit enlargement and experienced rotting after 25 days. Interestingly, when ABA and BR were applied to water lily stigmas treated with ISCP, it partially prevented the rotting of the ovary, significantly improving the fruit set rate and yielding a certain number of hybrid seeds. On the other hand, GA treatment, despite not promoting pollen tube germination on ISCP-treated stigmas, stimulated the enlargement of water lily ovaries, resulting in an increased fruit set rate. However, the majority of seeds in the fruits showed abortion, making it difficult to obtain viable seeds (Fig. 2CDE).

The germination experiment was conducted on the obtained seeds, it was found that the seeds treated with SP and CP exhibited a higher germination rate. Additionally, the hybrid seeds obtained through ISCP treatment with exogenously applied ABA and BR were also able to germinate successfully (Fig. 2F). However, compared to the plants obtained through SP and CP treatments, the hybrid seeds obtained through ISCP treatment required a longer germination time and exhibited a significantly slower plant growth rate (Fig. 2F).

### ABA and BR synergistically promote antioxidant enzyme activities in water lily stigmas

Reactive oxygen species (ROS) play a significant role in regulating the interaction between water lily pollen and the stigma, and the previous studies have also demonstrated this (Sun *et al*., 2023; Sun *et al*., 2019). Superoxide dismutase (SOD), peroxidase (POD), and catalase (CAT) are crucial antioxidant enzymes that directly regulate ROS accumulation (Kao *et al*., 2018). To gain further insights into how ABA and BR promote pollen germination on ISCP water lily, we assessed the activity of antioxidant enzymes in the stigma following pollination. Our results showed that the higher SOD, POD and CAT activity were observed in the stigma treated with SP and CP, while lower activity was observed in the Mock and ISCP stigmas. Furthermore, exogenous application of ABA and BR on the ISCP stigma promoted the enzyme activity of SOD, POD and CAT (Fig. 3CDE). Studies have shown that there is a synergistic regulation between ABA and BR (An *et al*., 2023; Li *et al*., 2021), in order to further understand the role of ABA and BR in the regulation of hybridization disorders between water lily subgenera. We further measured the levels of ABA and BR in the stigma. The results revealed that BR application on the ISCP stigma promoted the accumulation of endogenous ABA content, while ABA application on the ISCP stigma increased the endogenous BR content (Fig. 3AB). This suggests a potential synergistic regulation of ABA and BR in the process of hybridization in water lily.

**Fig. 3.**
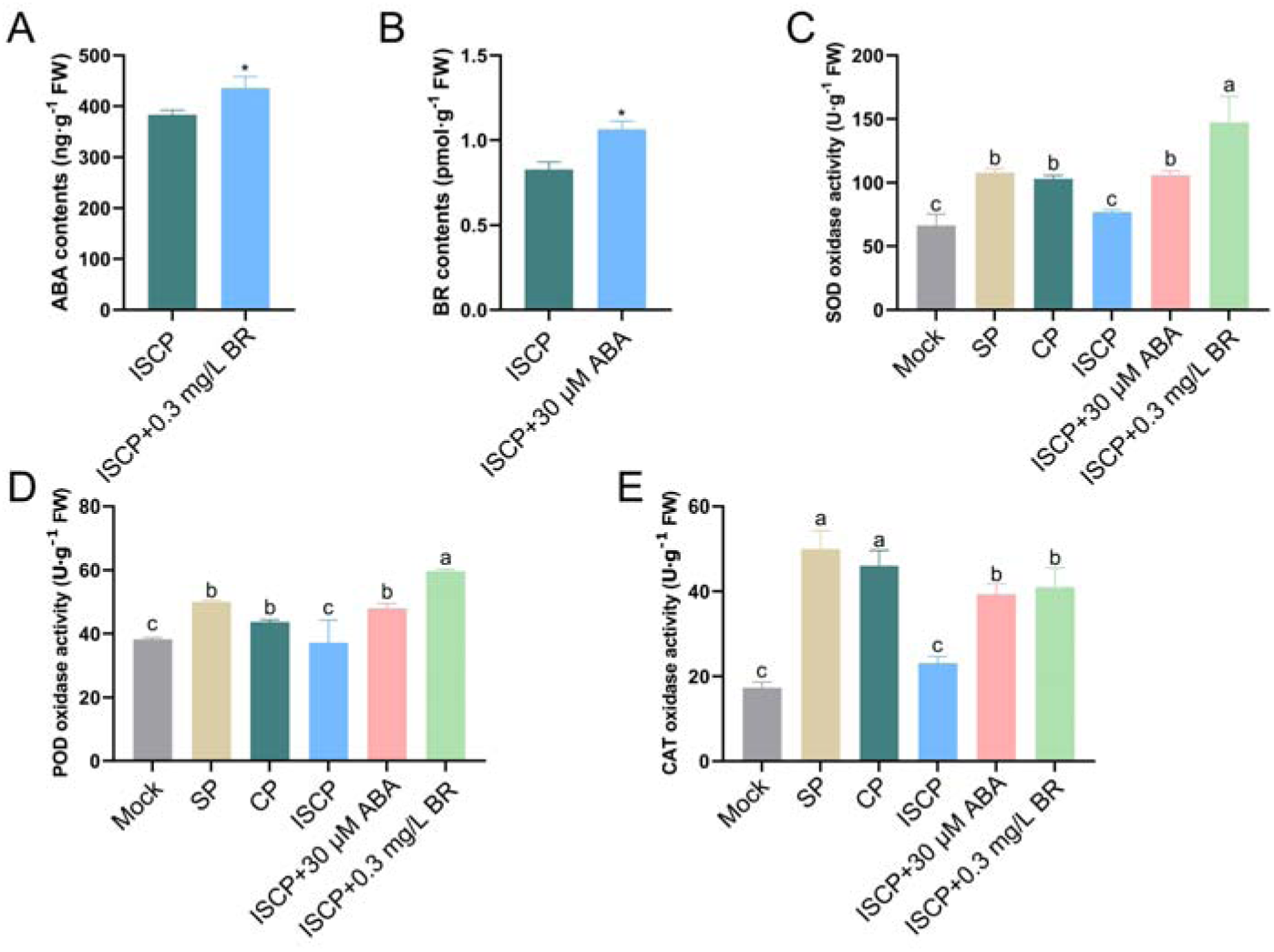
ABA and BR synergistically promote the activity of antioxidant enzymes in stigmas of water lily. (A) Determination of ABA content in water lily stigmas with ISCP or ISCP and BR treatment. (B) Determination of BR content in water lily stigmas with ISCP or ISCP and ABA treatment. (C-E) Activity assay of SOD, POD and CAT enzyme activity in water lily stigmas under different treatments. Different letters indicate statistically significant differences (*P* < *0.05*).

### Transcription profiles of water lily stigmas from different pollination combinations

In order to gain a deeper understanding of the specific mechanisms underlying the inter-subgeneric hybridization barriers in water lily, we utilized RNA-seq technology to investigate the gene transcription levels in stigmas treated with Mock, SP, CP and ISCP. The experimental design consisted of four sample groups, with four independent biological replicates for each sample, resulting in a total of 16 RNA-seq libraries. On average, each library generated 45.81 million single-end reads, ranging from 41.76 million to 50.44 million. The average number of clean reads was 43.92 million, ranging from 40.06 million to 49.56 million. Among these reads, 97.81% had a base quality score of Q20, and 93.65% had a base quality score of Q30 (Supplementary Table S2). Principal component analysis (PCA) based on FPKM values revealed that the first principal component (PC1) explained 40.8% of the variance in the dataset, while the second principal component (PC2) explained 25.3% of the variance. The Mock, SP, CP, and ISCP treatments formed distinct clusters, with the SP and CP groups exhibiting relatively closer clustering, and the clustering of all biological replicates was well-differentiated (Supplementary Fig. S2). These data indicate the reproducibility and reliability of the RNA-seq data.

Venn plot analysis revealed that in the ISCP_VS_Mock comparison, there were 1262 upregulated differentially expressed genes (DEGs) and 610 downregulated DEGs that were unique to this group (Fig. 4AB). These DEGs may play an important role in the inter-subgeneric hybridization barriers in water lily. To gain further insights into the functions of these 1872 DEGs, we conducted GO and KEGG analyses. The KEGG results showed that the 1262 upregulated genes were mainly involved in signal transduction, membrane transport, and cellular cytoskeleton (Fig. 4C). Additionally, the GO analysis indicated that the upregulated genes were mainly enriched in kinase activity and signal transduction, with Ca^2+^ transport and Ca^2+^ signaling exhibiting significant enrichment (Supplementary Fig. S3A). On the other hand, the 610 downregulated genes were primarily enriched in secondary metabolites, such as phenylpropanoid biosynthesis and flavonoid biosynthesis (Fig. 4D). The GO analysis revealed that the downregulated genes were mainly enriched in processes such as abscisic acid metabolism, carotenoid metabolism, and protein phosphorylation (Supplementary Fig. S3B).

**Fig. 4.**
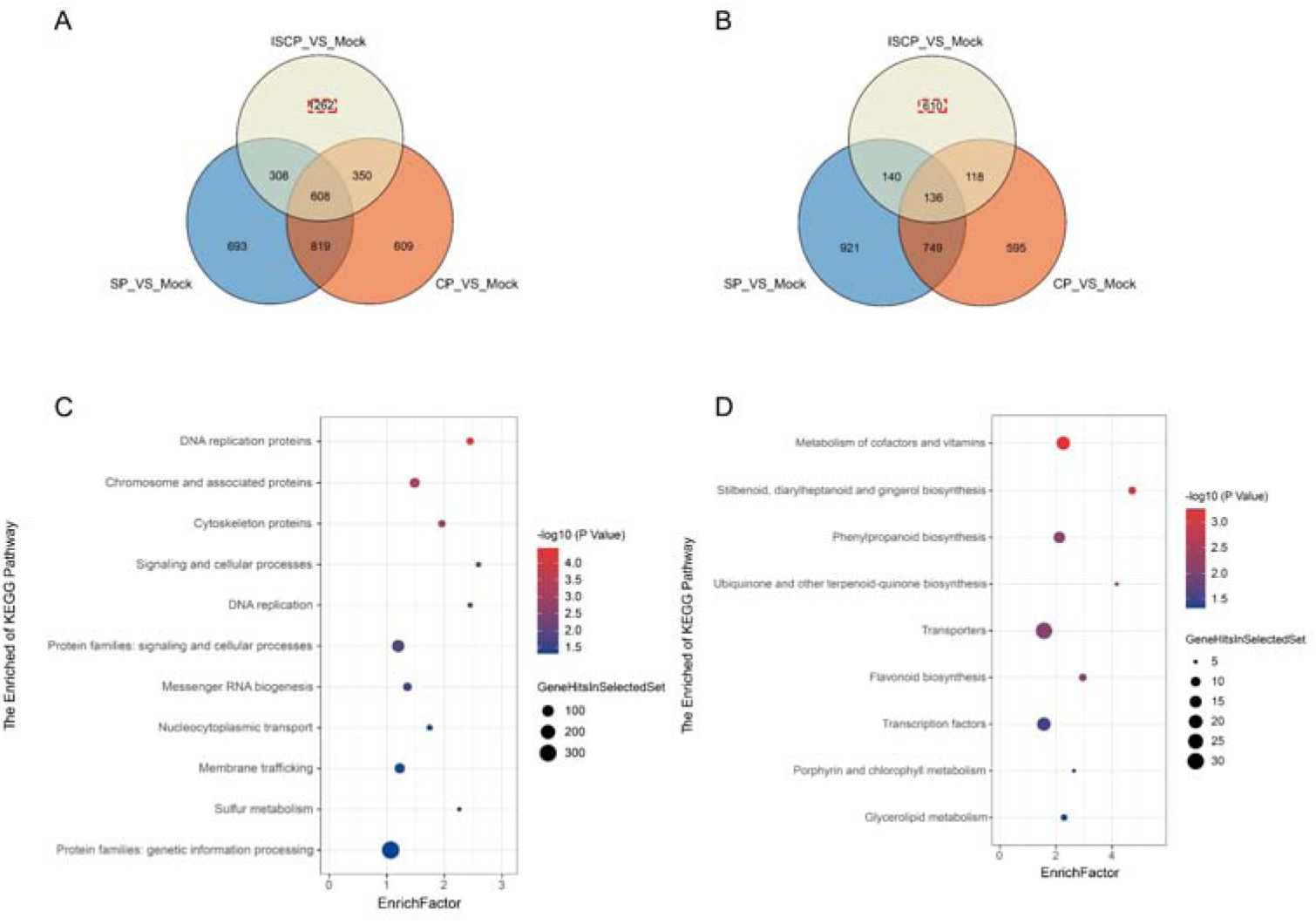
Identification of differentially expressed genes (DEGs) in ISCP stigma of water lily. (A-B) Venn diagrams showing the number of DEGs in comparison with Mock and the DEGs that are specifically up-regulated (A) or down-regulated (B) in the ISCP_VS_Mock grouping. (C-D) KEGG enrichment analysis of specifically up-regulated (C) or down-regulated (D) DEGs in the ISCP_VS_Mock grouping.

### Co-expression network analysis and key gene identification

The above results indicated that the changes of hormone levels play a significant role in regulating the formation of inter-subgeneric hybridization barriers in water lily. To understand the regulatory mechanisms behind the variations in hormone levels, we performed weighted gene co-expression network analysis (WGCNA) using the changes in ABA, GA3, MeJA, SAG and BR contents as phenotype data. The genes were divided into 26 co-expression modules using the WGCNA package (Supplementary Fig. S4 ABC). Genes within the same module exhibited highly similar expression patterns and were considered to be tightly co-regulated. The correlation between the phenotype data and the modules was visualized using the LinkET R package (Fig. 5A). The results showed that the magenta, turquoise, blue and yellow modules were positively correlated with MeJA and SAG contents, while the lightcyan module was negatively correlated with MeJA and SAG contents. Modules associated with ABA, GA3 and BR contents exhibited high similarity, such as the yellow, tan, black, brown and darkturquoise modules, which were positively correlated with these three hormones. On the other hand, the green, turquoise, blue and orange modules showed negative correlations with ABA and BR contents (Fig. 5A; Supplementary Fig. S4D).

**Fig. 5.**
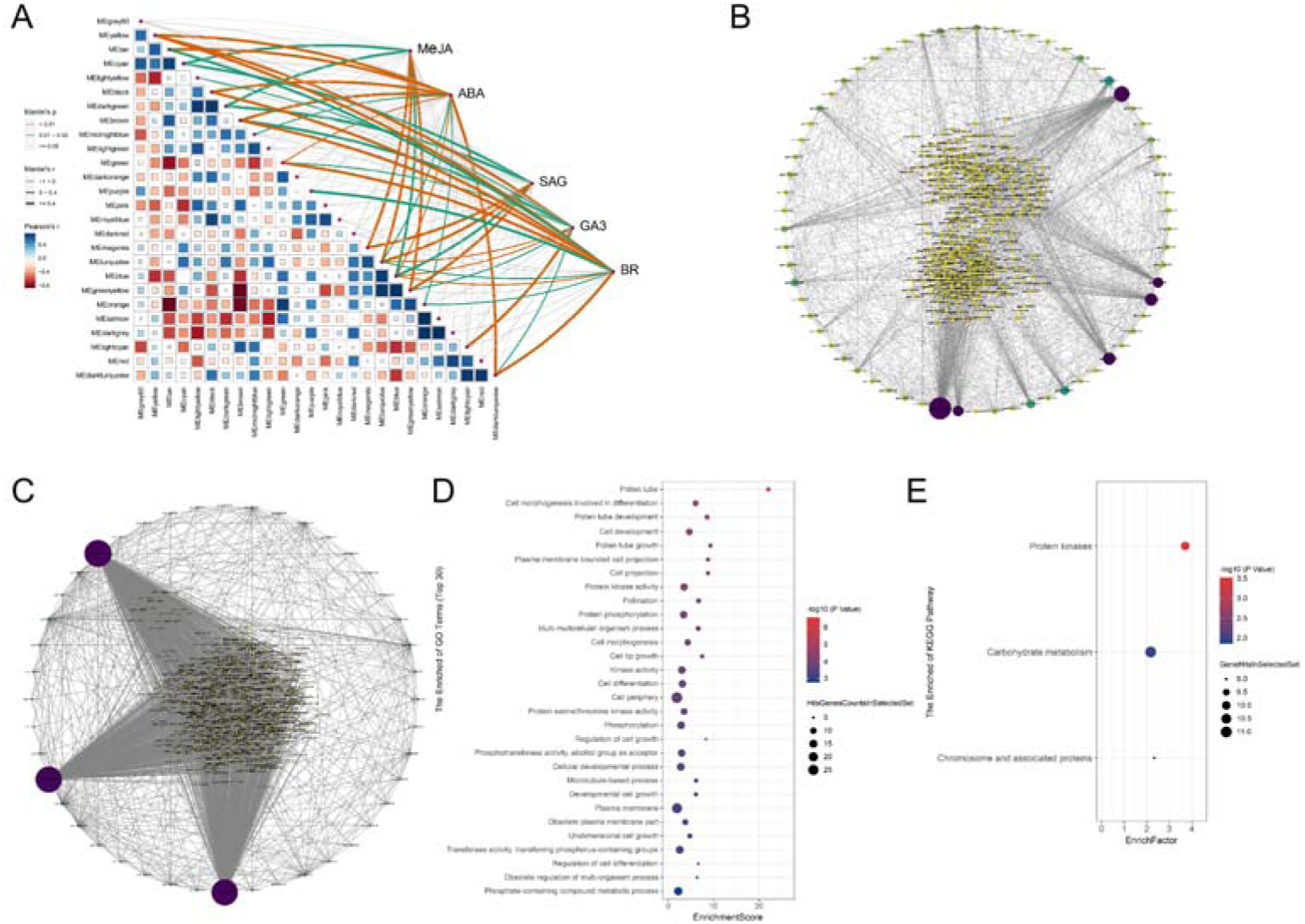
Identification of candidate genes for regulating inter-subgenus hybridization barriers in waterlily using weighted gene co-expression network analysis (WGCNA). (A) Module trait associations based on Pearson correlations. (B) Co-expression network of key genes from the blue module. (C) Co-expression network of key genes from the turquoise module. (D) GO enrichment analysis of key genes. (E) KEGG enrichment analysis of key genes.

It is interesting that two modules, blue and turquoise, showed correlations with all five hormones (Fig. 5A). Therefore, we selected the blue and turquoise modules for further analysis. GO and KEGG enrichment analysis revealed that genes in the blue module were mainly enriched in cellular cytoskeleton movement, signal transduction, kinase activity, and phosphorylation (Supplementary Fig. S5AC). On the other hand, genes in the turquoise module were primarily enriched in regulating pollen tube growth, organ development, cell growth and kinase activity (Fig. S5BD). We constructed a gene regulatory network using WGCNA and visualized the network using Cytoscape software (Fig. 5BC). We extracted the genes with the most connections, referred to as hub genes, and performed GO and KEGG analysis on them. The GO enrichment analysis showed that these hub genes were primarily enriched in regulating pollen tube growth, cell development, kinase activity, and phosphorylation (Fig. 5D). KEGG enrichment analysis revealed enrichment in protein kinases, carbohydrate metabolism, and chromatin-related proteins (Fig. 5E).

### Identification of genes involved in hormone biosynthesis

To further understand the relationship between the obtained hub-genes and the levels of these five hormones. We conducted an analysis of the expression of structural genes involved in the biosynthesis of these hormones (Fig. 6A) and performed a correlation analysis with the hub genes. The results revealed that key genes involved in ABA biosynthesis, named *ZEP*, *NCEDs*, *ABA2.1*, and *AAO3*, exhibited high expression levels in the stigmas of SP and CP treatments, which was consistent with the observed changes in ABA content. *KAO*, *GA3ox* and *GA20X1* were identified as potential important factors contributing to the accumulation of GA3 in the stigmas of SP and CP treatments. Within the BR synthesis pathway, most structural genes such as *CYP90A1* and *CYP85A1/2* were directly associated with the accumulation of BR in the stigmas of SP and CP treatments. The specific expression of *LOX* and *OPCL* in ISCP stigma is the main reason for the specific presence of MeJA in ISCP-treated stigmas. In the SA synthesis pathway, both the branched-chain pathway and the phenylalanine pathway appeared to participate in regulating the synthesis of SAG in ISCP-treated stigmas. Genes such as *ICS* in the branched-chain pathway and *PAL1.4* in the phenylalanine pathway were identified as potential important factors leading to the specific accumulation of SAG in the transmitting tract of the pistil.

**Fig. 6.**
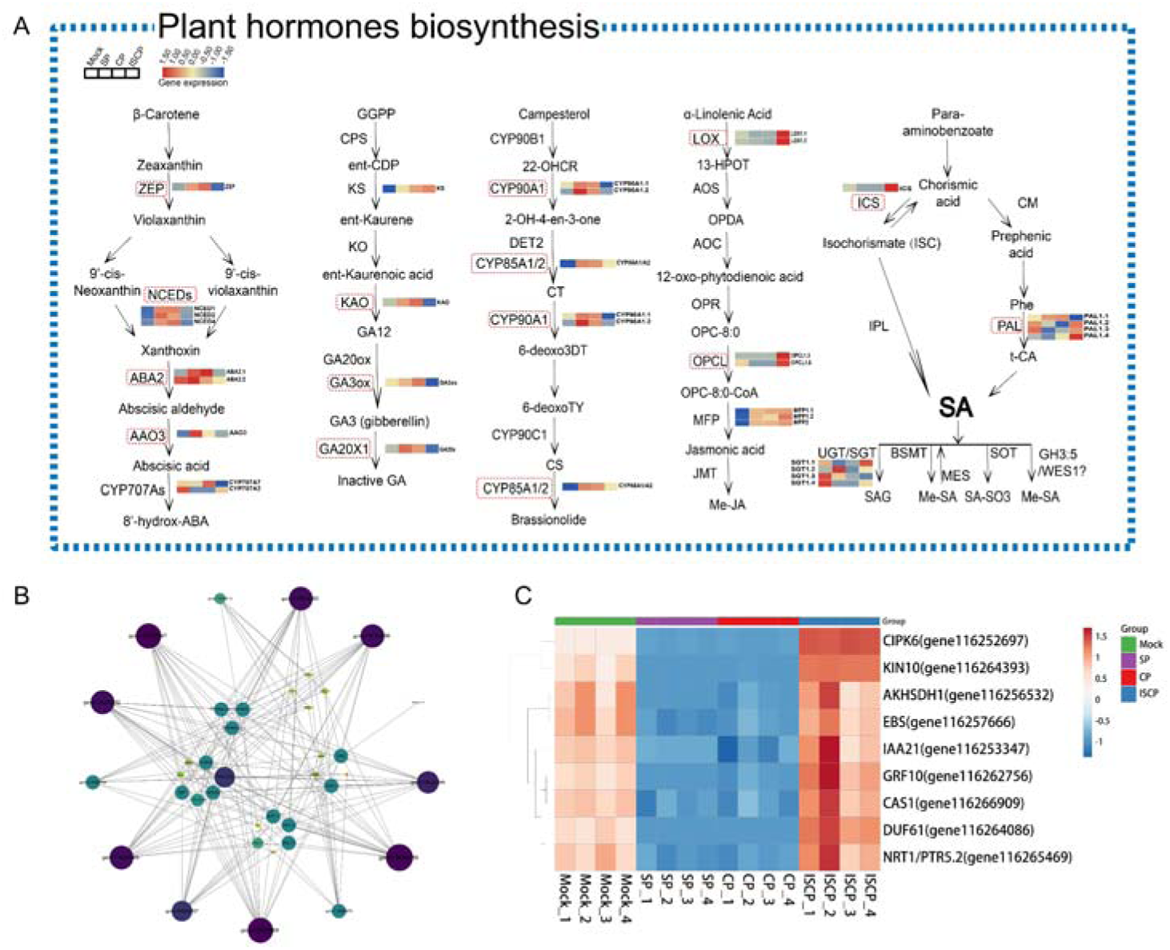
Correlation analysis of hormone biosynthesis genes with hub-genes. (A) Expression analysis of five hormone biosynthesis-related genes. (B) Correlation analysis of hub-gene with five hormone biosynthesis-related genes. (C) Heatmap showing the transcript levels of 9 hub-genes in Mock, SP, CP and ISCP treated stigmas (The transcript level information used to create the heat map is available in Supplemental Table S3).

The top 13 genes with the highest connectivity in the blue and turquoise modules (genes marked in purple and blue-green of Fig. 5B and genes marked in purple of Fig. 5C) were subjected to correlation analysis with the structural genes involved in the synthesis of the five hormones (Fig. 6B). The results revealed that the expression of nine genes was highly correlated with the expression of the structural genes involved in the synthesis of the five hormones. Additionally, we observed that *NCED2* had the highest number of connections among all the structural genes, suggesting that *NCED2* may be a key node gene for the regulation of these nine candidate genes. Visualization of the expression of these nine genes showed that their expression was low in the stigmas of SP and CP treatments, but high in the Mock and ISCP treatments (Fig. 6C).

In order to further validate the response of candidate genes to different pollination in water lily, qRT-PCR was used to confirm the expression of the candidate genes. The results showed that the expression of most genes was consistent with the transcriptome results (Fig. 7). Interestingly, among these structural genes, *NCED2*, *NCED4*, and *CYP707A2* may be responsible for the accumulation of ABA in the stigmas of SP and CP treatments. The changes in GA3 content may be attributed to the specific expression of *KAO* and *GA3ox* in the stigmas of SP and CP treatments. The genes mainly involved in BR and SAG synthesis were *CYP90A1*, *CYP85A1/A2*, *PAL1.2*, and *SGT1.1* (Fig. 7A). Among the selected nine candidate genes, all of them were highly expressed in the stigma of ISCP treatment. Among them, *CIPK6*, *KIN10*, *AKHSDH1*, *EBS*, *DUF61*, *NRT1/PTR5.2*, and *CAS1* may play more critical roles (Fig. 7B). The qRT-PCR results further suggest that the high expression of these nine genes in the stigma of ISCP treatment contributes to the formation of inter-subgeneric hybrid barrier in water lily.

**Fig. 7.**
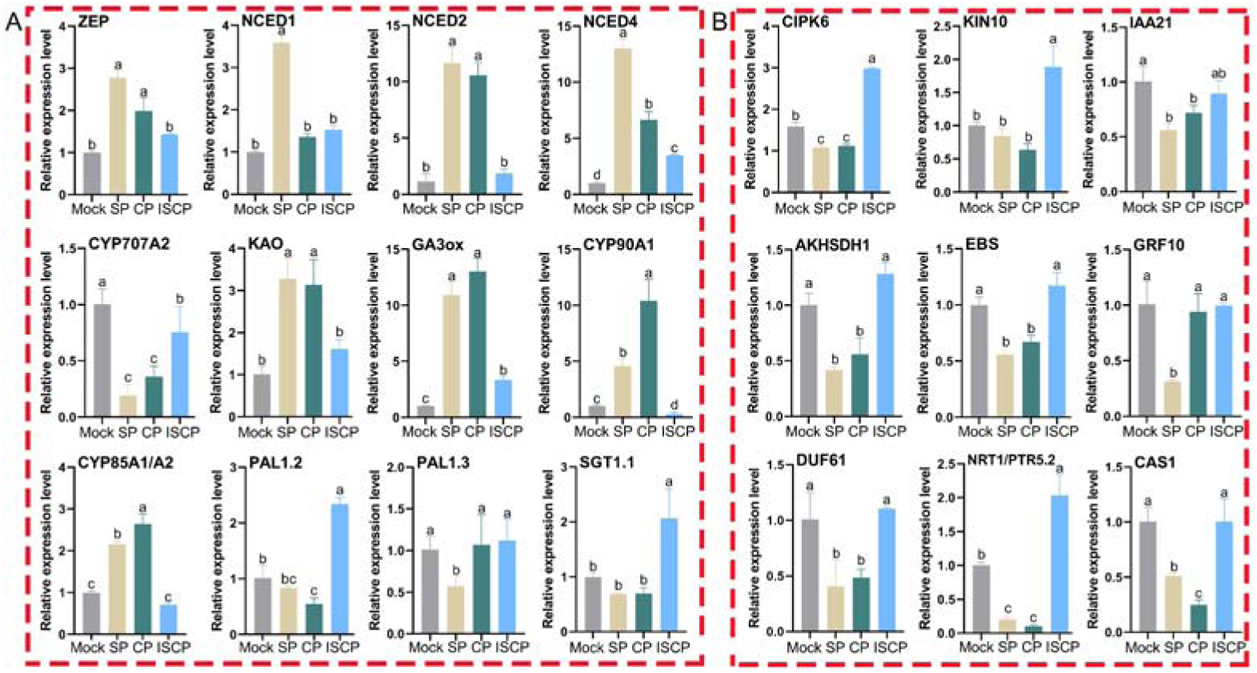
Validation of RNA-seq analysis of hormone biosynthesis and candidate genes in water lily. (A) qRT-PCR analysis of hormone biosynthesis genes. (B) qRT-PCR analysis of 9 candidate genes. All data were mean ± SD of three biological replicates, each containing three technical replicates. Different letters indicate statistically significant differences (*P* < *0.05*).

### Silencing of NCED2 on SP and CP stigmas inhibits pollen tube germination

The high connectivity of *NCED2* among all the structural genes in the network caught our interest. We further investigated the expression of *NCED2* in the stigmas of SP and CP treatments. The antisense oligonucleotide (ASO)-mediated was used to transient silence the expression of *NCED2*, we observed a significant reduction in the expression of *NCED2* in the stigma of SP and CP treatments (Fig. 8DE). The results showed that, when *NCED2* was silenced, the abundant pollen tube germination on the stigmas of SP and CP treatments were disappeared (Fig. 8ABC), accompanied by a decrease in ABA content (Fig. 8FG). However, when exogenous ABA was applied, pollen tube germination resumed on the stigmas (Fig. 8A). These results indicated that NCED2 is an important rate-limiting enzyme in ABA synthesis in the water lily pistil and that NCED2 promotes pollen germination by increasing ABA accumulation on the stigmas.

**Fig. 8.**
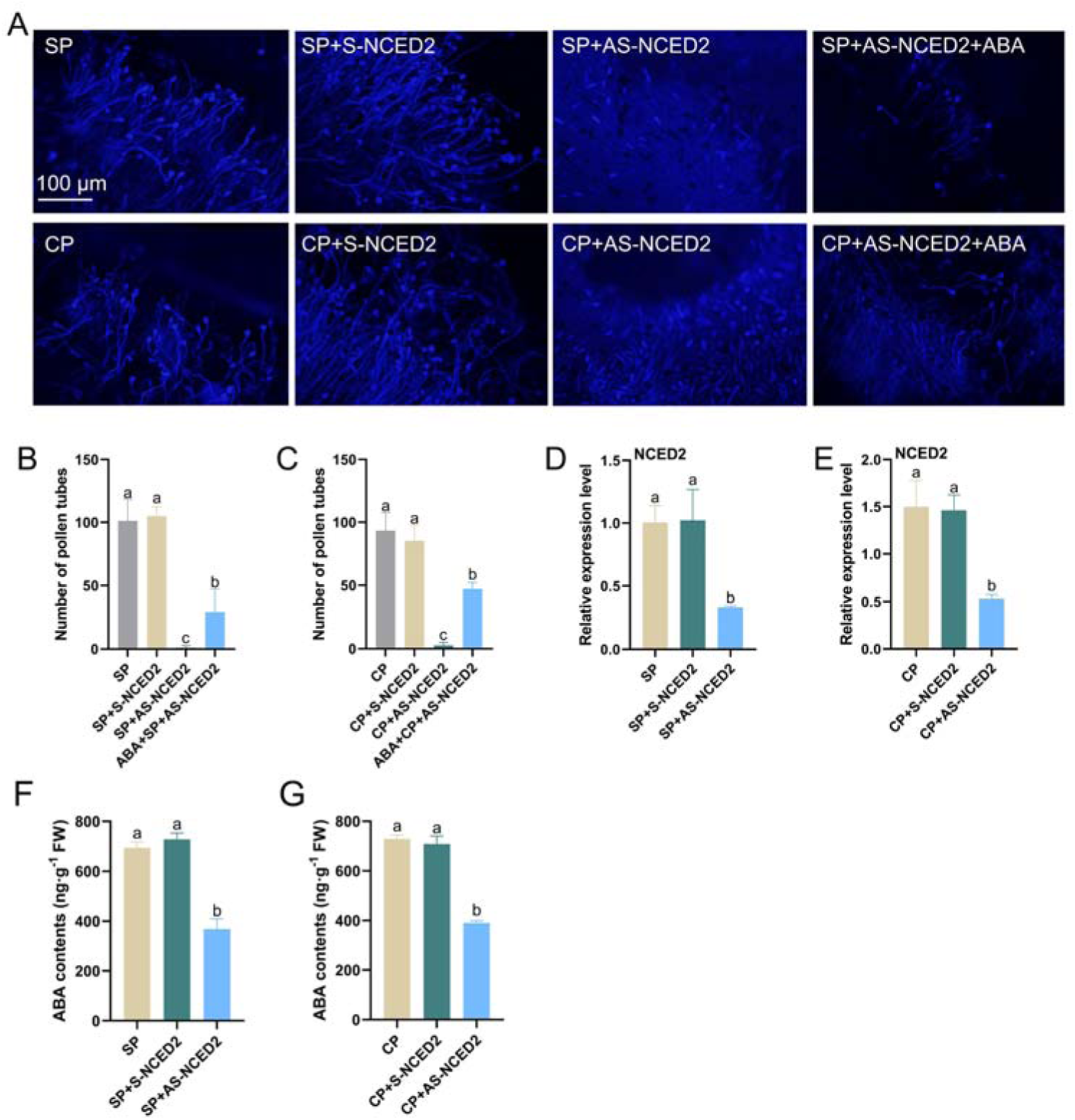
NCED2-mediated ABA synthesis may be a key to break the barrier of inter-subgenus in water lily. (A) Aniline blue staining showing the germination of pollen on water lily stigmas, Scale bar = 100 μm. (B-C) Statistics on the number of pollen germinating on water lily stigmas. (D-E) Transcript levels of NCED2 in differently treated stigmas. (F-G) ABA content assay after S or AS-NCED2 treatment of stigmas. All data were mean ± SD of three biological replicates, each containing three technical replicates. Different letters indicate statistically significant differences (*P* < *0.05*).

## Discussion

Plant hormones play important regulatory roles in the pollination and fertilization processes of plants, and research in this area has received widespread attention (Yang *et al*., 2022). Previous studies have shown that when the stigma receives compatible pollen, hormones related to growth and development significantly increase, such as auxin, gibberellins, and brassinosteroids (Qian *et al*., 2021; Shi *et al*., 2017; Wang *et al*., 2023; Wu *et al*., 2008; Ye *et al*., 2010). On the other hand, when the stigma receives incompatible pollen, hormones associated with plant stress tolerance significantly increase, such as ethylene, jasmonic acid and salicylic acid (Jegadeesan *et al*., 2018; Liu *et al*., 2022; Shi *et al*., 2017). In order to reveal the relationship between inter-subgeneric hybridization barriers and hormones in water lily, comprehensive transcriptomic and plant hormone analyses were conducted on the stigmas of water lily treated with Mock, SP, CP and ISCP. Additionally, we investigated the role of hormones in regulating inter-subgeneric hybridization barriers in water lily, with particular emphasis on the importance of ABA in in overcoming these barriers (Fig. 2; Fig. 8).

### The role of hormones in pollen-stigma communication of water lily

The germination and development of plant pollen are closely regulated by hormones (Calabrese and Agathokleous, 2021). IAA and ZR play a favorable role in the affinity between pollen and pistil (Yang *et al*., 2009), IAA playing an important role in promoting pollen tube growth (Wu *et al*., 2008). ZR, as the main natural active ingredient of cytokinins, is closely related to plant fertilization (Yang *et al*., 2022). Higher IAA, ZR levels and (IAA + ZR) / ABA levels favored pollination while lower ABA levels were detrimental during pollen-stigma recognition, adhesion and germination (Yang *et al*., 2022). In the process of hybrid breeding in water lily, the difficulty of pollen germination on the stigma is the main factor leading to inter-subgeneric hybrid barriers (Hao *et al*., 2022; Sun *et al*., 2023; Sun *et al*., 2019; Sun *et al*., 2018). In production, the spraying of exogenous hormones can be used to promote and increase the success rate of pollination, ensuring the progress of hybrid seed production (Daza *et al*., 2021).

Our results seemed to be consistent with the findings described above. Hormones associated with growth and development mainly accumulate in SP and CP stigmas, such as GA3 and BR. However, hormones accumulated in ISCP stigma are mainly related to biotic stress, such as MeJA and SAG (Fig. 1). Interestingly, in our results, ABA appears to positively regulate the affinity for water lily hybridization, mainly accumulating in SP and CP stigmas (Fig. 1). Studies have shown that maintaining a steady-state level of ABA is important for pollen germination and promoting pollen growth on the stigma (Lu *et al*., 2023). Higher levels of ABA hinder the normal development of the anther, thus reducing male fertility (Oliver *et al*., 2005; Oshino *et al*., 2007; Sharma and Nayyar, 2014). Increased levels of ABA in anthers lead to excessive accumulation of ROS, causing oxidative damage to anther tissues and disrupting the normal process of PCD in the chorioallantoic layer, which in turn affects the fertility of males (Sinha *et al*., 2021). While lower levels of ABA resulted in abnormal pollen development, thus affecting fertility (Kim *et al*., 2020; Lu *et al*., 2023). Endogenous ABA negatively affects pollen germination (Kovaleva *et al*., 2005), but exogenous ABA promotes pollen germination and pollen tube elongation in *P. hybrida* L self-compatible clones (Kovaleva *et al*., 2017). By exogenously applying ABA, GA and BR to ISCP stigmas, we found that both ABA and BR treatments can promote pollen germination on ISCP stigmas and facilitated the production of seeds through inter-subgeneric hybridization in water lily (Fig. 2). Further experiments demonstrated a synergistic regulatory effect between ABA and BR (Fig. 3), consistent with the results of (An *et al*., 2023; Li *et al*., 2021). They helped to alleviate ROS accumulation on ISCP stigmas by promoting the activities of antioxidant enzymes such as SOD, POD and CAT, thus breaking the inter-subgeneric hybridization barriers in water lily (Fig. 3).

### Signal transduction related genes and ROS play important roles in pollen-stigma communication of water lily

The process of pollen-pistil recognition involves complex signal communication, which is the first step for pollen germination on the stigma and ultimately leads to successful plant hybridization (Chapman and Goring, 2010; Jany *et al*., 2019; Johnson *et al*., 2019). To further understand the recognition mechanism between pollen and stigma during water lily hybridization and uncover the regulatory mechanisms involved in inter-subgeneric hybridization barriers in water lily, we conducted RNA-seq analysis of water lily stigmas with Mock, SP, CP and ISCP treatments. A total of 1262 genes specifically upregulated in the stigma of ISCP treatment and 610 genes specifically downregulated in the stigma of ISCP treatment were identified (Fig. 4AB).

Among the 1262 upregulated genes, their functions are mainly enriched in signal transduction pathways, such as Ca^2+^ transport, Ca^2+^ signal transduction and kinase activity. Conversely, the downregulated genes are primarily enriched in the synthesis of certain metabolites, including flavonoid synthesis, carotenoids and ABA synthesis (Fig. 4CD; Supplementary Fig. S3). Calcium (Ca^2+^) serves as a crucial regulator in mediating multiple signal transduction pathways involved in pollen germination and pollen tube growth (Nie *et al*., 2023). In pear hybridization, disruption of cytoplasmic Ca^2+^ oscillations in incompatible pollen and disturbance of the focused Ca^2+^ gradient at the tip of developing pollen tubes severely impairs polar growth of the pollen tubes (Jiang *et al*., 2014; Qu *et al*., 2016). High levels of Ca^2+^ can induce programmed cell death events in incompatible pollen on the stigma of *Papaveraceae* (Wilkins *et al*., 2014). In *Brassicaceae*, the self-incompatibility response triggered an immediate increase in Ca^2+^ concentration within the stigma, promoting extensive callose deposition, thereby suppressing the germination and pollen tube elongation of incompatible pollen (Iwano *et al*., 2015). These findings suggested a negative regulatory role for Ca^2+^ in the communication process between pollen and the stigma. Interestingly, in our findings, genes associated with flavonoid synthesis, carotenoid synthesis, ABA synthesis and other related pathways showed a downregulation, and these metabolites are involved in the clearance of reactive oxygen species (ROS).

Previous studies have suggested that the high accumulation of ROS in the stigma of water lily is an important factor contributing to reproductive barriers in inter-subgeneric hybrids (Sun *et al*., 2023; Sun *et al*., 2019). Furthermore, elevated ROS levels have also been shown to promote self-incompatibility and cross-incompatibility in other species. For example, in *Brassicaceae*, the levels of ROS in the stigma play a role in regulating both self-incompatibility and interspecies hybrid barriers (Huang *et al*., 2023; Zhang *et al*., 2021). In *Arabidopsis*, variations in ROS levels in the stigma directly impact pollen hydration and germination on the transmitting tract cells (Liu *et al*., 2021). Therefore, we propose that the accumulation of ROS mediated by Ca^2+^ signaling in the stigma of water lily may be one of the key factors contributing to the formation of reproductive barriers in inter-subgeneric hybrids. Further WGCNA analysis was performed and two modules, blue and turquoise, were screened from 26 expression modules (Supplementary Fig. S4). These two modules showed significant correlation with the content of MeJA, SAG, ABA, GA3 and BR (Fig. 5A). Nine candidate genes (*CIPK6, KIN10, IAA21, AKHSDH1, EBS, DUF61, NRT1/PTR5.2* and *CAS1*) were finally obtained by correlation analysis. The functions of these nine genes were involved in Ca^2+^ signaling (*CIPK6*), AUX response (*IAA21*) and hormone transporter-related (*NRT1/PTR5.2*). Furthermore, these results provide further evidence for the significance of Ca^2+^ and hormone signaling in the formation of hybridization barriers in waterlily.

### NCED2-mediated ABA synthesis is a critical for promoting pollen germination on water lily stigmas

NCED is a key rate-limiting enzyme in ABA synthesis that catalyzes the oxidative cleavage of 9-cis-violaxanthin or 9-cis-neoxanthin, producing xanthoxin in the plastids (Gupta *et al*., 2022). Study have shown that RNAi-mediated silencing of the *NCED* gene reduces ABA levels and inhibits pollen development in transgenic tomatoes (Dai *et al*., 2018). Interestingly, overexpression of *NCED* also suppresses pollen development, despite the increase in ABA levels, indicating that pollen development is sensitive to ABA homeostasis (Dai *et al*., 2018; Wang *et al*., 2021b). In our results, the reduction in ABA levels in the stigma of water lily (stigma of ISCP treatment) is an important factor contributing to reproductive barriers in inter-subgeneric hybrids. Additionally, NCED2 plays a crucial role in ABA synthesis in the stigma of water lily (Fig. 6AB; Fig. 7A). To investigate the regulatory role of NCED2 in reproductive barriers in water lily, we treated the stigmas of SP and CP with antisense oligodeoxynucleotides (AS-ODNs). The results showed that AS-ODN treatment significantly reduced the expression of *NCED2* in the stigmas of SP and CP (Fig. 8DE). Moreover, silencing of *NCED2* hindered pollen germination on the stigma, while ABA treatment restored pollen germination (Fig. 8A).

In conclusion, we have found that hormones play an important role in regulating reproductive barriers in inter-subgeneric hybrids of water lily. Among these hormones, ABA and BR may have a more critical role and show potential in overcoming reproductive barriers in water lily. RNA-seq analysis has revealed the involvement of Ca^2+^ signaling and some metabolites with antioxidant capacity in regulating reproductive barriers. Based on these results, we have proposed a working model (Fig. 9). After receiving pollen, the stigma of water lily activates Ca^2+^ signaling and regulates pollen germination and growth by modulating the synthesis of ABA, BR and some metabolites such as flavonoids and carotenoids. Our results provide guidance for improving hybridization efficiency and overcoming reproductive barriers in water lily. It also provides a theoretical basis for further understanding of the mechanisms that regulate hybridization barriers in inter-subgeneric hybrids of water lily.

**Fig. 9.**
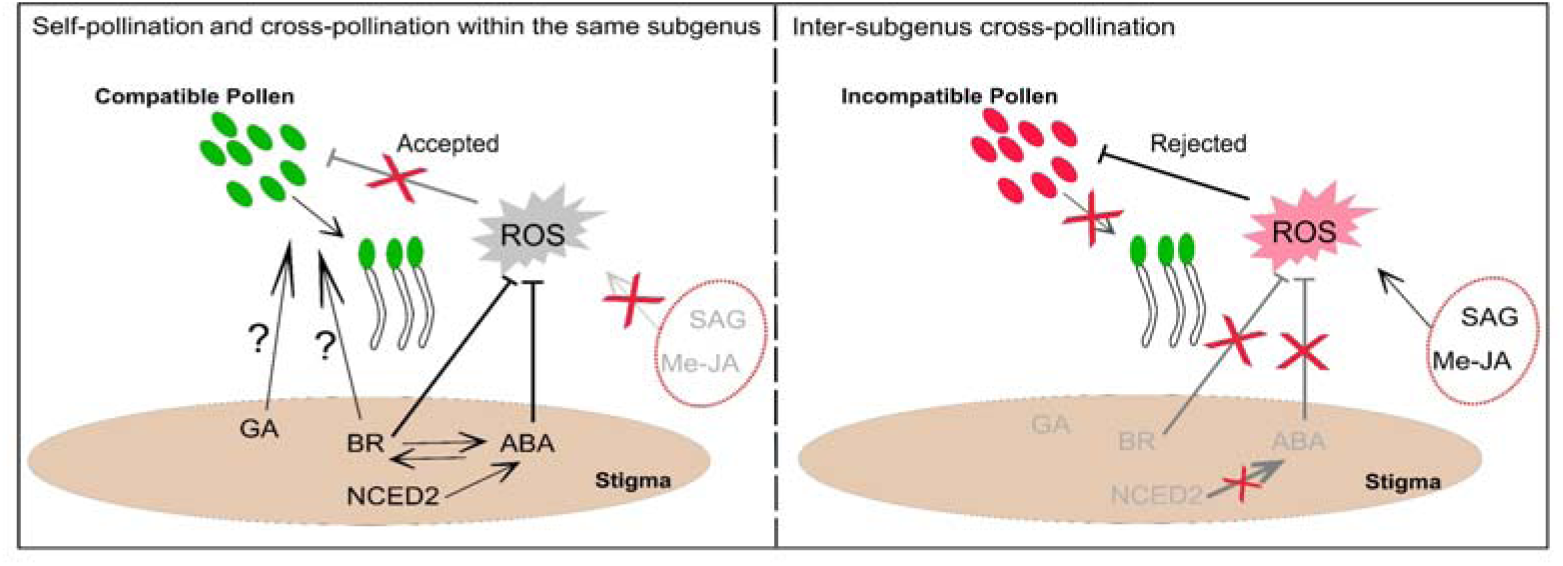
The model of hormones in regulating hybridization the barriers of inter-subgenus in water lily.

## Acknowledgements

We would like to thank Suzhou Panomics Biomedical Technology Co., Ltd (Suzhou, China) for providing hormone and transcriptome testing services.

## Author Contributions

PZ and YCX conceived the original idea and designed the experiments; PZ, JWL, HYJ, ZJY and HYW performed the experiments; YJW provided the experimental materials; PZ analyzed the data and wrote the manuscript; CQS, QS and QJJ revised the manuscript. All authors have read and approved the manuscript.

## Data Availability

The RNA-seq data were submitted to the BIG Data Center (http://bigd.big.ac.cn) under the BioProject accession PRJCA018998.

## The following Supporting Information is available for this article

Fig. S1. Changes in the abundance of phytohormones.

Fig. S2. Principal component analysis of the 16 samples used for transcriptomics analysis.

Fig. S3. GO enrichment analysis of unique DEGs in the ISCP_VS_Mock grouping.

Fig. S4. Identification of key genes associated with MeJA, ABA, SAG, GA3 and BR content by WGCNA analysis. (A) Co-expression modules were identified using a hierarchical clustering tree. 26 identified modules are indicated with different colors. (B) Heatmap showing the topological overlap matrix of DEGs. Fading colors indicate lower overlap and red indicates higher overlap. (C) Correlation analysis between 26 co-expression modules. (D) Correlation analysis between co-expression modules and the content of five hormones.

Fig. S5. GO and KEGG enrichment analysis of genes in blue and turquoise modules. (A) GO enrichment analysis of genes in the blue module. (B) GO enrichment analysis of genes in the turquoise module. (C) KEGG enrichment analysis of genes in the blue module. (D) KEGG enrichment analysis of genes in the turquoise module.

Table S1. Detection of hormone content in water lily stigmas.

Table S2. Summary of RNA-Seq data.

Table S3. Gene expression information from the transcriptome used in the heatmap.

Table S4. Primers used for qRT-PCR analysis.

Table S5. S or AS-ODN sequences.

